# Event-related network changes unfold the dynamics of cortical integration during face processing

**DOI:** 10.1101/2020.06.29.177436

**Authors:** Antonio Maffei, Paola Sessa

**Affiliations:** Padova Neuroscience Center (PNC), University of Padova, Italy; Department of Developmental and Social Psychology, University of Padova, Italy

**Keywords:** Time-varying connectivity, Face processing, EEG, Dynamic networks, Brain modularity, Small-worldness

## Abstract

Face perception arises from a collective activation of brain regions in the occipital, parietal and temporal cortices. Despite wide acknowledgement that these regions act in an intertwined network, the network behavior itself is poorly understood. Here we present a study in which time-varying connectivity estimated from EEG activity elicited by facial expressions presentation was characterized using graph-theoretical measures of node centrality and global network topology. Results revealed that face perception results from a dynamic reshaping of the network architecture, characterized by the emergence of hubs located in the occipital and temporal regions of the scalp. The importance of these nodes can be observed from early stages of visual processing and reaches a climax in the same time-window in which the face-sensitive N170 is observed. Furthermore, using Granger causality, we found that the time-evolving centrality of these nodes is associated with ERP amplitude, providing a direct link between the network state and local neural response. Additionally, investigating global network topology by means of small-worldness and modularity, we found that face processing requires a functional network with a strong small-world organization that maximizes integration, at the cost of segregated subdivisions. Interestingly, we found that this architecture is not static, but instead it is implemented by the network from stimulus onset to ~200 msec. Altogether, this study reveals the event-related changes underlying face processing at the network level, suggesting that a distributed processing mechanism operates through dynamically weighting the contribution of the cortical regions involved.

**Data Availability:** Data and code related to this manuscript can be accessed through the OSF at this link https://osf.io/hc3sk/?view_only=af52bc4295c044ffbbd3be019cc083f4

## 1. Introduction

When presented with a face, the human brain reacts with a fast cascade of highly integrated responses that result in the perception of the stimulus as a face, and in the extraction of the large amount of information a face can convey, like identity, gender, age, emotional expression, attractiveness and so forth (Adolphs & Birmingham, 2011; Dobs, Isik, Pantazis, & Kanwisher, 2019). Decades of research into the neural basis of human face perception established that this ability results from an activity distributed across several brain areas, comprising altogether a face perception network (Grill-Spector, Weiner, Kay, & Gomez, 2017; Haxby & Gobbini, 2012; Haxby, Hoffman, & Gobbini, 2000; Ishai, 2008; Nguyen, Breakspear, & Cunnington, 2014; Tsao & Livingstone, 2008). This network comprises occipito-temporal regions involved in processing the basic visual features comprising a face, as well as, in building an holistic percept of these features (the “core” system of the network), but also parietal and fronto-central regions involved in coding high-level features like face identity and emotional information conveyed by facial expressions (the “extended” system of the network) (Grill-Spector et al., 2017; Haxby & Gobbini, 2011). Despite the acknowledgment that face processing relies on the activity of a set of functionally interconnected regions, rather than a single brain area, the behavior of this network is still poorly understood. This is especially true with regard to the time-evolving dynamics that govern network activity (Dobs et al., 2019). Indeed, functional MRI has an inherent limited time resolution that allows modelling the interaction of brain regions in this network in a static fashion (Fairhall & Ishai, 2007) or, at best, on a slower time scale compared to other techniques (Preti, Bolton, & Van De Ville, 2017), but prevents to achieve a fine understanding of how the network activity changes rapidly over time. (Magneto)Electroencephalography, on the other hand, has an excellent time-resolution and has been widely used to characterize with exceptional detail the fast time evolution of face processing. Using non-invasive (magneto-)electrophysiological techniques has been possible to finely measure brain activity subtending this unique ability, uncovering that faces prompt a positive voltage peak around 100 msec (the P1) followed by a negative voltage deflection peaking at 170 msec (the N170). This N170 activity recorded from the posterior regions of the scalp represents a hallmark of face processing in the brain, and it is widely accepted that these evoked response represents the endpoint of the mazy mechanism underlying the perception of “faceness” (Eimer, 2012; Itier & Taylor, 2004; Rossion, 2014). In terms of event-related oscillations (ERD/ERS), face processing is associated with an increased power in gamma band (Rossion, 2014) mediated by an increased synchronization in slow frequency oscillations, especially in the alpha band (Rousselet, Husk, Bennett, & Sekuler, 2007; Yang, Qiu, & Schouten, 2015). Furthermore, changes in the alpha power has been linked with emotional expressions and the expressivity conveyed by facial movements (Girges, Wright, Spencer, & O’Brien, 2014; Güntekin & Başar, 2014). Unfortunately, investigating face processing only by means of ERPs and ERD/ERS can inform us just on the endpoint of this mechanism, but cannot provide direct information on the dynamics within the network that eventually result in this typical neural response.

Integrating EEG with fMRI-based connectivity modeling has proven useful to characterize the dynamics of face perception in terms of regional cross-communication patterns (Nguyen et al., 2014), showing that increased N170 amplitude is associated with an increased effective connectivity between the regions belonging to the *core* system. To further improve our understanding of these dynamics, is nevertheless necessary to focus on electrophysiological estimates of connectivity, which can inform us about how these dynamics unfold with an excellent time-resolution (Valencia, Martinerie, Dupont, & Chavez, 2008). Indeed, previous studies that investigated connectivity with EEG showed an increased synchronization between cortical regions, especially in the alpha band (Yang et al., 2015). Furthermore, they showed that this synchronization occurs mostly between posterior and anterior regions, in line with the predictions of a distributed systems (Yang et al., 2015).

The present study aimed at providing new insights about these dynamics, leveraging the power of graph theory applied on time-resolved connectivity estimated from EEG activity elicited by the presentation of facial expressions, contrasted with the activity elicited by non-face stimuli (i.e., animal shapes). The innovation of this study lies in the combination of graph-theoretical indices of nodal importance and global network topology with an advanced non-parametric statistical framework, to characterize the event-related network topological changes associated with face processing. In line with the growing field of functional *chronnectomics* (Calhoun, Miller, Pearlson, & Adali, 2014; Preti et al., 2017), by studying the evolution of the network we aimed to eventually increase current knowledge regarding the timings of the neural states subtending cognitive mechanisms of face processing.

For what concerns the characterization of each node activation state, we first investigated what the network hubs are (i.e. the most important nodes in the network) and how their importance evolves in time. Second, we asked if the knowledge of the state of the network is associated with the typical face-evoked ERPs. To answer these questions, we identified network hubs by means of three different graph-theoretical measures of centrality, namely degree, PageRank and eigenvector centrality (Zuo et al., 2012). We advanced a strong prediction that the major hubs (i.e. the node showing highest centrality during face processing) will correspond to the occipito-temporal sensors from which the N170 response is typically observed with largest amplitude. Additionally, we expected that their importance will progressively increase from stimulus onset, peaking around 200 msec (i.e. the time window in which face-related ERPs show the largest amplitude). Finally, we employed Granger causality to test the relationship between centrality time-series and the ERPs.

For what concerns the topological properties of the whole network, we asked if the topology of the network changes depending on the stimulus under processing (face vs. non-face) and how these changes can inform on the time-evolution of the neural computations within the network.

Although there is no doubt that face processing involves a collection of regions, there is an ongoing debate on how this network operates. This debate is characterized by two, partly competitive, alternatives suggesting that the face network relies on either a modular or a distributed information processing mechanism. According to the former hypothesis, within the face network there should be specific brain areas showing face-selective activity (Kanwisher & Barton, 2011; Kanwisher & Yovel, 2006) that operate in segregated modules and along anatomically and functionally separated dorsal and ventral pathways (Duchaine & Yovel, 2015; Freiwald, Duchaine, & Yovel, 2016). According to the latter hypothesis, face processing can be viewed instead as an emergent property of the activity of a distributed information processing mechanism that does not necessary rely on face-specific circuits, but operates on domain-general circuits (Behrmann & Plaut, 2013; Haxby & Gobbini, 2012; Schrouff et al., 2020). In support to a dynamic distributed mechanism, several experimental evidence also suggests that the face network response is shaped by stimulus features and task characteristics (Fairhall & Ishai, 2007; Mechelli, Price, Friston, & Ishai, 2004; Summerfield et al., 2006). Specifically, the link between the core and the extended systems appears to be selectively modulated by these features. Presenting participants with emotional facial expressions increases the connectivity between the fusiform gyrus and the amygdala, while presenting famous and attractive faces increases the connectivity between the fusiform and the orbitofrontal cortex (Fairhall & Ishai, 2007). Moreover, the ability to judge if a stimulus is a face or not is related to feedback projections that functionally connect the ventral frontal cortex to the core system (Summerfield et al., 2006).

To answer these questions in order to gain a better understanding of the principles governing the information exchange within the cortical face network, we focused on two important topological properties that brain networks possess, namely small-world topology and modularity. Small-Worldness refers to the property of many real-life systems characterized by having a topology with a high local clustering (i.e. neighbor nodes are highly interconnected) and, at the same time, a short average path length among nodes (i.e. average number of connections between a random pair of nodes is low) (Watts & Strogatz, 1998). Such property entails for biological networks, and hence for the brain, the potential for supporting information processing that can be both segregated and distributed (Bassett & Bullmore, 2006; Telesford, Joyce, Hayasaka, Burdette, & Laurienti, 2011). On the other hand, modularity defines the tendency of network nodes to segregate into distinct modules characterized by strong within-modules connectivity and low between-modules connectivity (Sporns & Betzel, 2016). Brain networks, both structural and functional, have been demonstrated to show a modular topology (Sporns & Betzel, 2016; van den Heuvel & Sporns, 2013), and recent evidence suggests that brain modularity is not static, but instead changes over time (Betzel, Fukushima, He, Zuo, & Sporns, 2016; Fukushima et al., 2018), as well as a function of the task (resting state vs. active condition) (Di, Gohel, Kim, & Biswal, 2013). These changes are believed to index the dynamic fluctuation between states of segregated (high modularity) and states of integrated (low modularity) information processing.

According to both theoretical perspectives on face processing we could predict that face stimuli will shape brain connectivity toward an increased clustering within a global small-world architecture, but modularity is expected to increase only if the processing depends on segregated modules. On the contrary, network modularity is predicted to decrease if a distributed processing mechanism is involved.

## 2. Methods

### 2.1. Participants

Thirty-four university students (6 males, mean age = 22.8 y, sd = 3.3 y) volunteered to take part to this research. All participants had normal or corrected-to-normal vision and reported no history of neurological or psychiatric diseases. The study was conducted according to the principles of the Declaration of Helsinki and after approval of the IRB of the University of Padova. All participants granted their written informed consent to participate in the study.

### 2.2. Stimuli and procedures

Stimuli were 11 grayscale digital photographs portraying animals (a cow or a horse) or faces displaying a negative emotional expression (anger or sadness) (Niedenthal, Halberstadt, Margolin, & Innes-Ker, 2000). For each face photograph, a set of morphed expressions was generated on a continuum starting from 100% sadness and 0% anger and ending to 0% sadness and 100% anger, in step of 10%. To provide a balanced control, the same morphing was adopted for the animal condition, using a continuum starting from 100 % cow and 0 % horse and ending to 0 % cow to 100 % horse. This control condition was designed to match the visual demands required by recognition of subtle facial expressions, retaining as much as possible the ecology of the visual experience of the participant and minimizing the confounds related to having different levels of difficulty between the experimental conditions. Using faces with emotional expressions, rather than neutral expressions, we sought to maximize the recruitment of the whole face perception network which is known to show an increased response to this kind of stimuli (Haxby & Gobbini, 2012; Hinojosa, Mercado, & Carretié, 2015). Images were resized to subtend a visual angle between 10° and 12°, and presented at the center of a computer screen placed 60 cm away from the participant.

Trials were structured in a 500 ms baseline period in which a fixation cross was displayed at the center of the screen, followed by the presentation of the target stimulus that lasted for 750 ms. After a 350 ms masking interval, the target image was presented along with a distractor image drafted from the corresponding continuum of morphed images. The task was to correctly identify the target image from the distractor. Inter-trial interval varied randomly between 800 and 900 msec. The total number of trials was 288.

**Figure 1:**
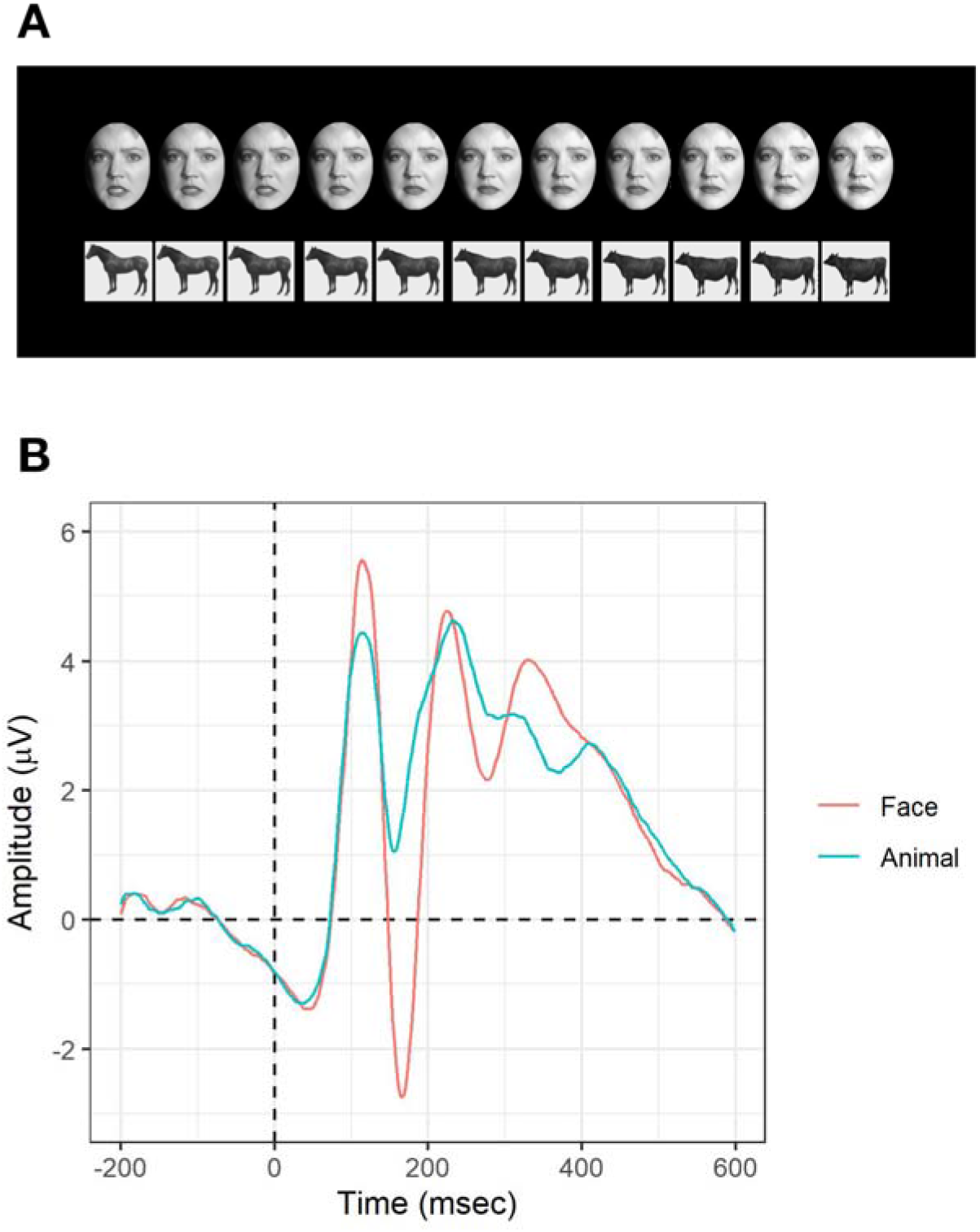
Morphed stimuli for face and animal categories (panel A) and their evoked response measured at sites PO7/PO8 (panel B)

### 2.3. EEG recording and preprocessing

Scalp electrical activity was collected using a BrainAmp amplifier, from 62 active electrodes placed on an elastic Acti-Cap according to the extended 10/20 system. Signals were recorded using the left ear lobe as online reference, with a sampling rate of 1000 Hz and keeping the electrodes impedance below 10 KΩ.

Continuous data were downsampled to 500 Hz, high-pass filtered at 0.1 Hz, re-referenced to the average of all channels and segmented in epochs starting from −500 ms to 750 ms with respect to stimulus onset. Independent component analysis (ICA) was applied to the epoched data in order to identify and discard artifacts related to eye-blinks and saccades (Jung et al., 2000).

Time-resolved connectivity between scalp sensors was derived computing, for each time point, the instantaneous phase locking value (Lachaux, Rodriguez, Martinerie, & Varela, 1999; Sakkalis, 2011) between each pair of sensors after transforming voltage activity into current source density (CSD) by taking the surface Laplacian of scalp voltage (Kayser & Tenke, 2015; Srinivasan, Winter, Ding, & Nunez, 2007). CSD transform is a necessary prerequisite to investigate functional connectivity at scalp level, since it provides a reference-free estimate of brain activity and significantly reduces the impact of volume conduction, thus improving the spatial resolution of surface EEG (Kayser & Tenke, 2015).

After CSD transform, the signals were first bandpass filtered in the alpha (8-13 Hz) band, then their analytical representation was extracted using the Hillbert transform and the phase angle differences between each sensors pair was computed for each time point, according to the formula:

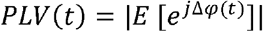

where Δ*φ*(*t*) represents the relative phase angles between two analytical signals *z*_1_(*t*) and *z*_2_(*t*) at each time point *t*. This analytical strategy resulted in a time-series of weighted undirected connectivity matrices that represented the basis for the computation of graph theoretical metrics, an approach that has been successfully applied also to characterize time-resolved connectivity from fMRI data (Nobukawa, Kikuchi, & Takahashi, 2019; Pedersen, Omidvarnia, Zalesky, & Jackson, 2018).

Due to the lack of previous works investigating functional connectivity during face processing with EEG, we focused on alpha band because of the evidence showing a strong overlap between functional connectivity estimated from BOLD response and functional connectivity estimated from alpha oscillations (Mantini et al., 2007). This choice aimed at providing insights which might be generalized across imaging modalities. Indeed, alpha oscillatory activity is strongly related to the activity of large-scale brain networks (Sadaghiani et al., 2010) and its phase synchrony has been shown to mediate the functional integration between cortical regions located to a relative distance from each other (Maffei, 2020; Muller, Chavane, Reynolds, & Sejnowski, 2018; van Driel, Knapen, van Es, & Cohen, 2014). Slower oscillations (< 10 Hz) are instead spread on the whole cortex and thus can inform only on integration mechanism that occur at very long time scales (Maffei, 2020; Massimini, Huber, Ferrarelli, Hill, & Tononi, 2004; Steriade, McCormick, & Sejnowski, 1993). Faster oscillations in the beta/theta range have been successfully targeted to study region-specific activity related to face processing, due to the very focal spread of these oscillations (Rossion, 2014). This very narrow spread has nonetheless a downside. Investigating functional connectivity in these spectral ranges might be less suited to inform on the communication and integration among distant regions (Maffei, 2020). On the other hand, the phase synchrony in the alpha range is appropriate for the purpose of this study, because of its role in coordinating gamma oscillations (Bahramisharif et al., 2013) related to complex sensory processing (Jerbi et al., 2009) and face processing (Rossion, 2014).

In order to compute ERPs, artifact-free data were further segmented into shorter epochs starting from −200 ms to 600 ms with respect to stimulus onset, baseline corrected and low pass filtered at 30 Hz. Epochs with a peak-to-peak amplitude exceeding ± 50 μV in any channel were identified using a moving window procedure (window size = 200 ms, step size = 50 ms) and discarded from the averaging procedure. Accordingly, all subsequent analyses on connectivity data were performed in the same time frame.

### 2.4. Graph-theory analyses

Graph-theory is a powerful mathematical theory that allows to model brain connectivity and to describe the characteristics of complex brain networks which rapidly became the gold-standard to study large scale brain connectivity (Bullmore & Sporns, 2009) and their time-evolving dynamics (Calhoun et al., 2014).

In this study, characterization of the importance of nodes within the functional network was performed combining three different indices of node centrality, namely degree centrality (DC), eigenvector centrality (EC) and PageRank centrality (PC). The use of three metrics, rather than a single one, stemmed from the need to find convergent evidence and to avoid drawing biased conclusion since each metric emphasizes different features of topological centrality (Zuo et al., 2012).

Node degree is one of the simplest centrality index and is derived by simply counting the number of edges connecting a node with all the others. Eigenvector centrality is instead defined for each node as the corresponding eigenvector related to the largest eigenvalue of the adjacency matrix, and is able to capture the influence of the global network topology on each node centrality score. The principle subtending EC is that not all connections have the same importance, thus influent nodes will be the ones connecting prominently with other influential nodes rather than with nodes with low centrality. Finally, PageRank centrality is computed from the well-known Google’s algorithm, and is variant of EC that assigns a damping factor *d* (*d* = 0.85, in this study) to control the likelihood that a random walker traveling the graph ends up in a sink. Although originally developed to find central nodes in directed graphs, PageRank algorithm has been successfully applied also to undirected graphs, showing a strong relationship to degree centrality (Zuo et al., 2012).

In addition to nodal measures, we computed two indices of global network topology, namely small-worldness and modularity. Small-worldness aims at describing the network topology by computing the ratio between its clustering coefficient and its path length (Watts & Strogatz, 1998). The higher this ratio the more the network has a “small-world” topology. In this study small-worldness (ω) was computed according to the following equation, originally proposed by Telesford and colleagues (2011):

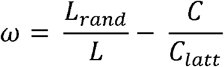

where *L* and *C* are, respectively, the average path length and the average clustering coefficient of the network, and *L*_rand_ and *C*_latt_ are, respectively, the average path length of a random network and the average clustering coefficient of a lattice network, both generated preserving the size and degree distribution of the original network.

The normalization procedure bounds the possible values of ω between −1 and 1, with a small world topology corresponding to values close to 0 (Telesford et al., 2011). Compared to the seminal Watts and Strogatz’s (1998) definition, this procedure deals more efficiently with the problem of normalizing the index by taking into account both a random and a lattice rewiring of the observed network, which represent the two extreme ends on the continuum that small-worldness aims at characterizing.

Modularity (*Q*) is a statistics that quantifies the optimal partitioning of a network into nonoverlapping subdivisions characterized by a large number of within-module edges and sparse between-modules connectivity (Newman, 2006). A common problem encountered with the study of networks modularity, is that finding the optimal subdivision can be a computationally intensive problem, thus optimization algorithms are normally used. Here, the Louvain “greedy” algorithm was employed to identify the community structure maximizing *Q* (Blondel, Guillaume, Lambiotte, & Lefebvre, 2008). Thus, the resulting modularity index shows how strong is the modular topology within the network, with higher values of *Q* suggesting a more clearly defined modular organization (Stanley, Dagenbach, Lyday, Burdette, & Laurienti, 2014).

Since the aim of the present work was to characterize the time-evolving dynamics of brain connectivity, all these graph-theoretical measures were computed from the adjacency matrix *A*_(*i*)_ reflecting the phase-coupling between nodes at each time-point *i*. This approach resulted in having, for each metric, a time series reflecting how that property unfolds over time. Furthermore, in order to deal with the problem of appropriately thresholding the adjacency matrix to ensure sufficient sparsity to avoid spurious connections, the observed connectivity matrices were thresholded using three different thresholds *ρ* = [.2 .1 .05], retaining respectively the 20%, 10% and 5% of the strongest connection weights. This procedure was applied for all the metrics. Finally, all the metrics were computed for the connectivity matrices estimated during face and animal trials.

The analyses were performed in MATLAB (v2019a) using custom scripts employing functions from the EEGLab toolbox (Delorme & Makeig, 2004), ERPLab toolbox (Lopez-Calderon & Luck, 2014), Brainstorm (Tadel, Baillet, Mosher, Pantazis, & Leahy, 2011) and Brain Connectivity Toolbox (Rubinov & Sporns, 2010).

### 2.5. Statistical Analysis

The statistical modeling of the time-varying graph properties was performed within a massive univariate non-parametric permutation framework (Groppe, Urbach, & Kutas, 2011). This framework consists in performing a statistical test (like a t-test or ANOVA) for every point in the electrode by time plane, then iteratively permuting the data and performing the test a sufficient number of times to have an empirical null-distribution of the test statistic from which derive the exact probability and then perform the statistical inference.

Non-parametric analysis based on permutations, combined with a cluster-based approach to handle the problem of multiple comparisons (Bullmore et al., 1999), represents the gold-standard for EEG/ERP analysis (Maris & Oostenveld, 2007). This approach allows to relax the very strict assumptions of parametric models and allows to take into account the full multidimensional structure of psychophysiological datasets without restricting *a-priori* the testing to specific set of electrodes and time-window(s).

Since the graph-theoretical metrics in this study were structured in a node by time format, we extended this framework in order to model the event-related topological changes in the network and to test at which node and in which time-window face stimuli prompted an increased centrality compared to animal stimuli.

This test was performed separately for each nodal property (DC, EC, PC) and threshold level of connectivity matrices (20%, 10%, 5%) using 5000 permutations performed taking into account the whole epoch duration (−200-600 msec). Statistical significance was assessed using α = 0.05 and employing a cluster-based approach for the control of multiple comparisons using the Fieldtrip’s *ft_ timelockstatistic* implemented in Brainstorm.

For the analysis of the two global topological metrics, i.e. small-worldness and modularity, the same approach was implemented in R (v 3.5.3) using 1000 permutations, α = 0.05 and handling the problem of multiple comparisons by controlling the false discovery rate with the Benjamini and Hochberg’s procedure (1995).

Finally, we tested the predictive value of nodal centrality scores over ERP amplitude elicited by face stimuli using Granger Causality. Given two time-series *x*[t] and *y*[t], Granger causality quantifies how the unique information in one of the time series improves the prediction of the values of the other. This is done by testing the hypothesis that including information of past values of *x* provides a better prediction of future values of *y*, compared to the prediction of *y* derived by its simple autoregressive function (Roebroeck, Formisano, & Goebel, 2005).

Using a univariate approach, we tested at each cortical site the causal influence (in the Granger sense) of each nodal centrality metric time-series over the corresponding ERP time-series. This was accomplished with an F-test comparing the ERP autoregressive model incorporating the appropriately lagged values of centrality to the ERP autoregressive model without this information. The appropriate MVAR lag was defined as the one minimizing to the Akaike Information Criterion (Matias et al., 2014). The opposite test (ERP time series predicting the centrality time-series) was also performed. Performing the test in both directions, allowed us to identify only nodes showing a significant (α = .05) Granger-causality of centrality on ERP and a non-significant Granger-causality of ERP over centrality, helping us to rule out ambiguous effects.

## 3. Results

### 3.1 Node Centrality

The analyses of the time-varying dynamics of nodes centrality showed a strong effect, consistently observed across different metrics and threshold, revealing a cluster of temporo-parieto-occipital nodes which are the most important during face processing (Figure 2). The analyses also revealed that their centrality becomes relevant in discriminating faces from non-face stimuli very early during stimulus processing and peaks around 200 msec from stimulus onset (Figure 3).

**Figure 2:**
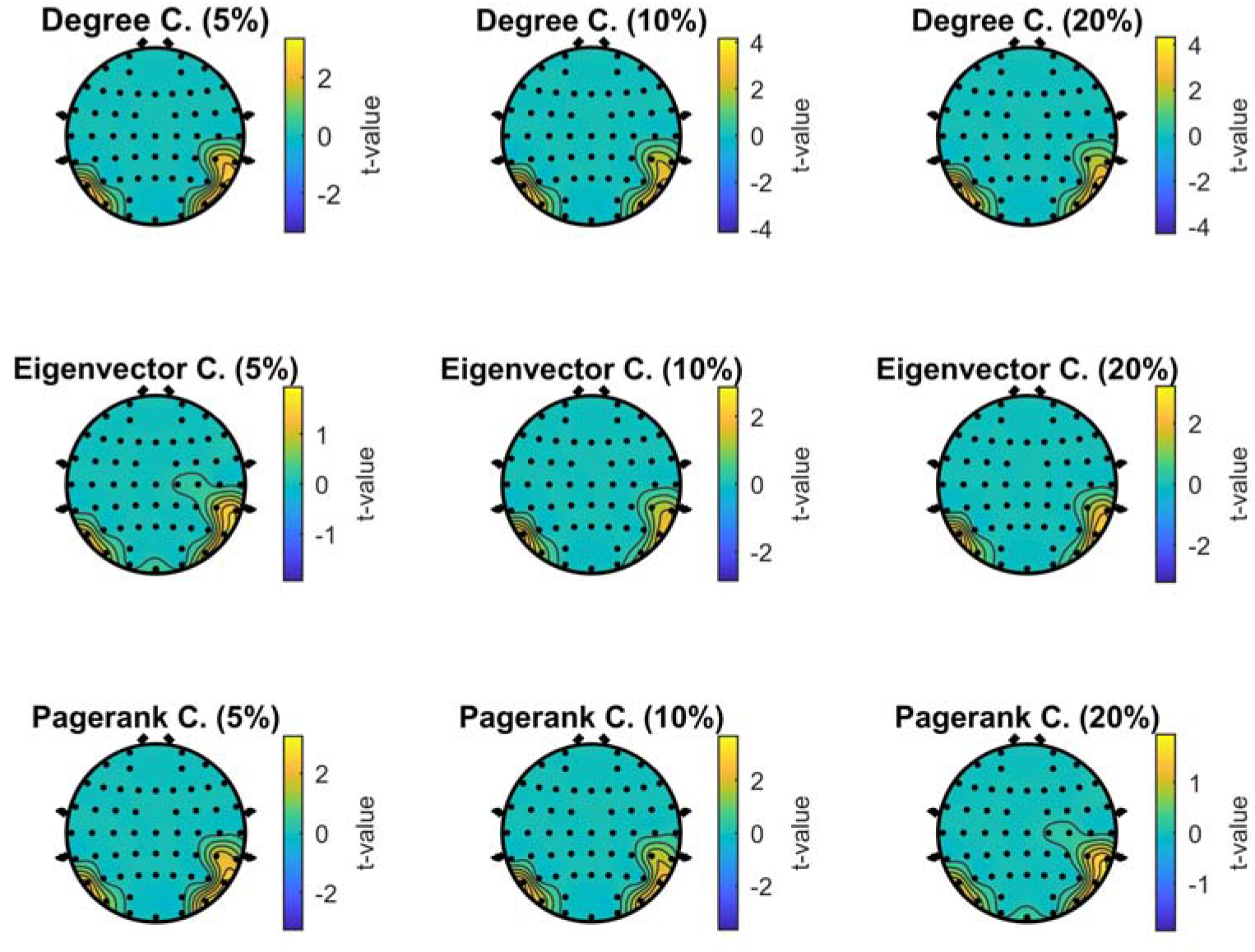
Scalp maps showing the significant statistical effect for the contrast between Face and Animal stimuli for each centrality metric and threshold

**Figure 3:**
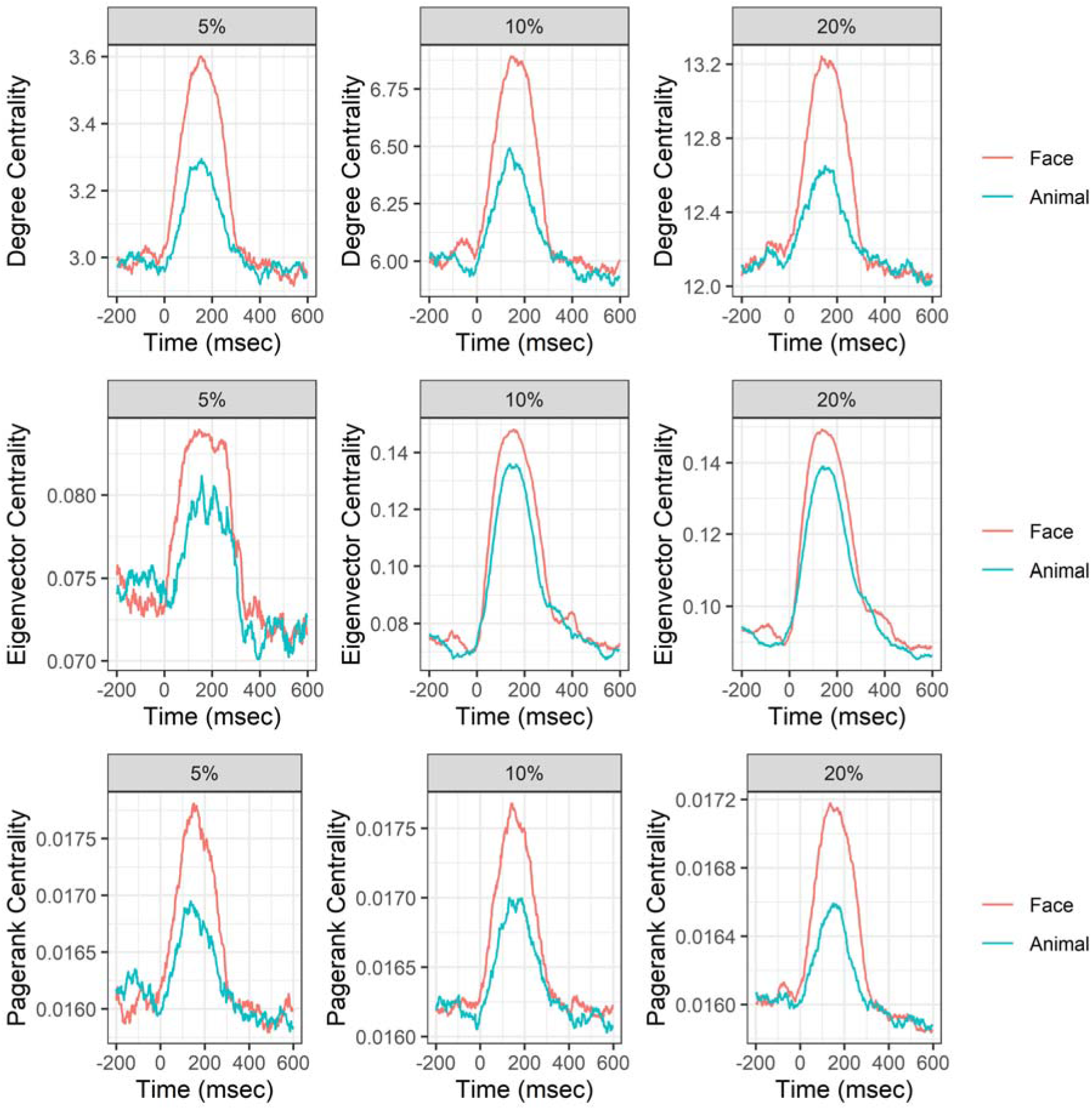
Time-series of the average centrality of the nodes showing a significant effect for the contrast between Face and Animal stimuli, for each centrality metric and threshold.

With regard to degree centrality (DC), cluster-based permutation tests revealed an increased centrality during face processing compared to animal processing for all the three thresholds considered (5 %: *t*_max_ = 2954, *p*-value < 0.0001, cluster size = 958; 10 %: *t*_max_ = 3230, *p*-value < 0.0001, clustersize = 956; 20 %: *t*_max_ = 3155, *p*-value < 0.0001, clustersize = 929).

Similar results were observed for cluster-based permutation tests performed on eigenvector centrality (EC), showing a strong difference between face and animal conditions for all thresholds (5 %: *t*_max_ = 1833, *p*-value = 0.009, cluster size = 741; 10 %: *t*_max_ = 2007, *p*-value = 0.012, clustersize = 781; 20 %: *t*_max_ = 2290, *p*-value = 0.011, clustersize = 822).

Finally, the same pattern was observed also for the cluster-based permutation tests performed on PageRank centrality scores (PC), revealing again a strong effect of face over animal processing for each threshold (5 %: *t*_max_ = 2755, *p*-value < 0.0001, cluster size = 912; 10 %: *t*_max_ = 3013, *p*-value < 0.0001, clustersize = 952; 20 %: *t*_max_ = 3136, *p*-value < 0.0001, clustersize = 928).

### 3.2 Granger-causality of graph properties on ERP amplitude

The analysis of the predictive power of node centrality over ERPs revealed significant effects, especially at medium level of thresholding (ρ = .1), in the right temporo-parietal regions of the scalp for both Degree centrality and PageRank centrality (Figure 4).

**Figure 4:**
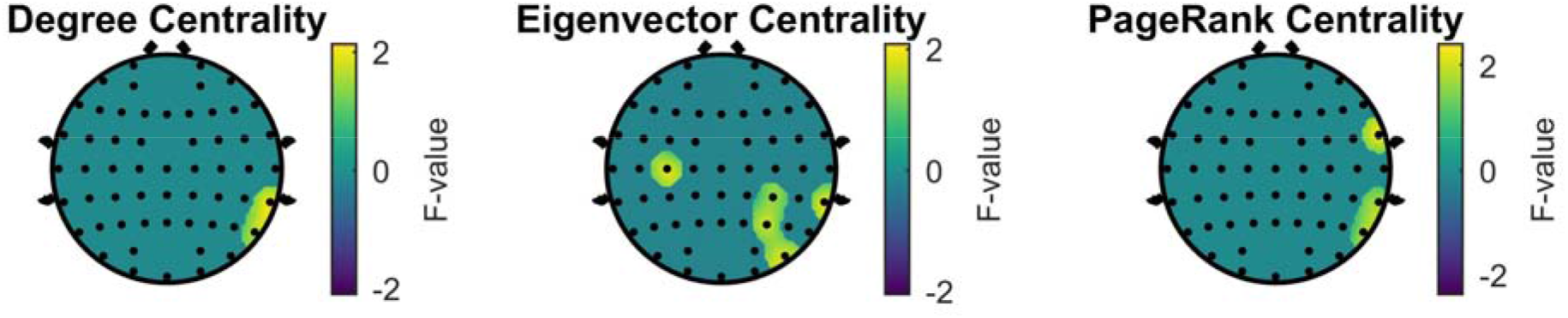
Scalp maps showing, for each metric, nodes in which was observed a significant effect in the Granger causality test of the predictive value of centrality on ERP amplitude.

DC showed a significant effect for the Granger-causality test at nodes P8 (F_(15,710)_ = 1.71, p = 0.04) and TP8 (F_(11,734)_ = 2, p = 0.02). PC showed a significant effect at nodes P8 (F_(15,710)_ = 1.7, p = 0.04), TP8 (F_(14,716)_ = 1.76, p = 0.03) and FT8 (F_(27,638)_ = 2.29, p < 0.0002). These effects were observed also using a stricter level of thresholding (p = .05), while using a more liberal thresholding (ρ = .2) the results showed an effect spreading over the cortex (see Supplementary material, for a list of significant nodes).

A similar pattern was observed also in the analysis performed on Eigenvector centrality (Figure 2), which further confirmed the predictive value of the centrality computed in nodes PO8 (F_(26,644)_ = 1.82, p = 0.007), TP8 (F_(13,722)_ = 1.94, p = 0.02) and P4 (F_(23,662)_ = 1.74, p = 0.01), additionally showing a predictive effect over central sites C3 (F_(15,710)_ = 1.86, p = 0.02) and CP4 (F_(34,596)_ = 1.45, p = 0.04).

The analyses performed using a liberal threshold (ρ = .2) showed instead an effect spreading over the cortex, while only in one node an effect was observed using a stricter thresholding (p = .05).

### 3.3 Global Topology

The analyses of the time-evolving properties of the global network topology revealed that face processing is supported by a small-world network that relies on a distributed processing organization. The analysis of the small-worldness (Figure 5A) showed a significant decrease of the parameter ω, approaching 0 which indexes a greater small-world topology (Telesford et al., 2011) during presentation of face stimuli compared to animal stimuli (5 %: *t*_max_ = 115, *p*-value < 0.05; 10 %: *t*_max_ = 497, *p*-value < 0.05; 20 %: *t*_max_ = 581, *p*-value < 0.05).

**Figure 5:**
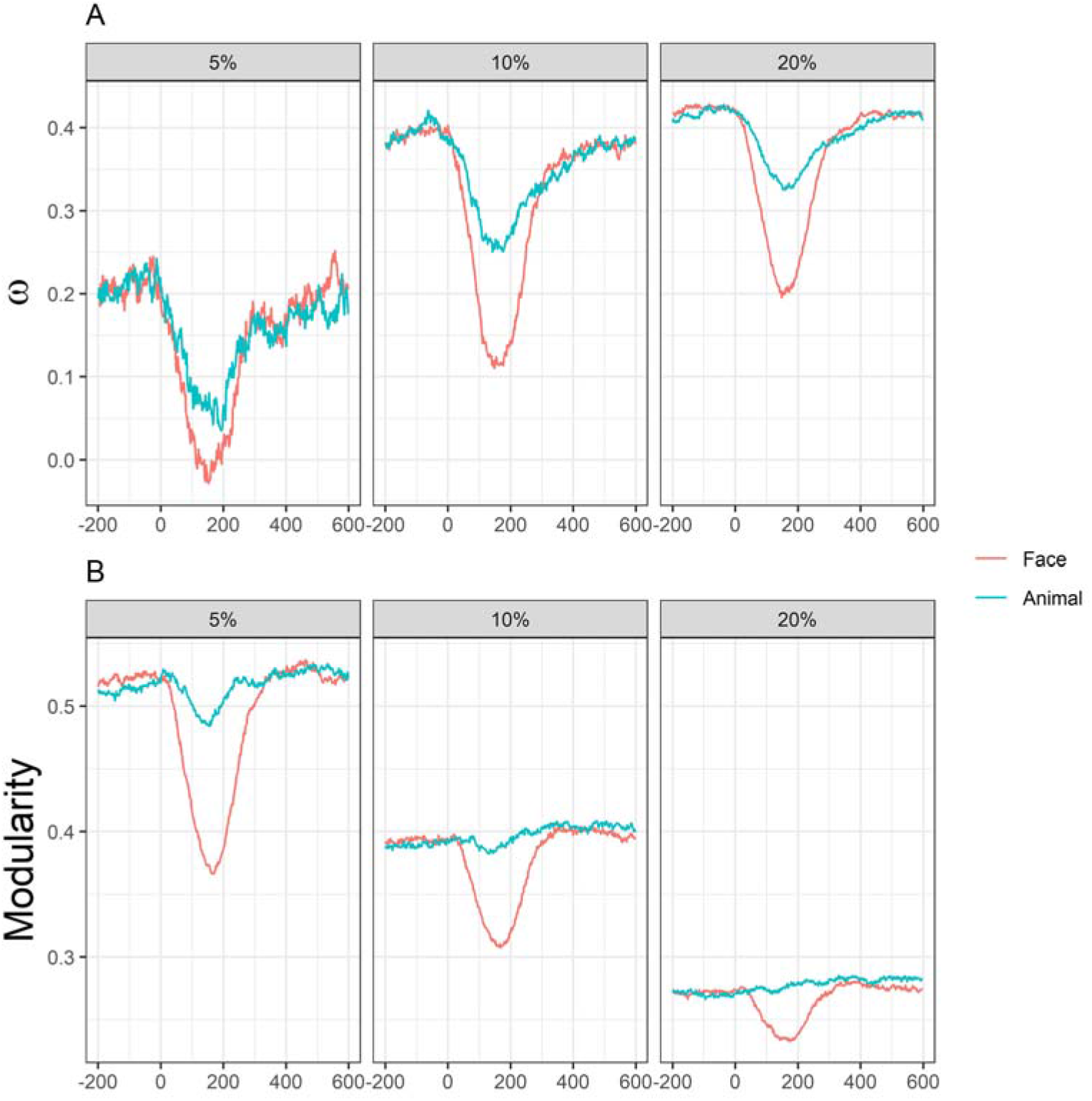
Top panel (A) shows the time-varying small-worldness index (ω) for Face and Animal stimuli observed at each threshold. Bottom panel (B) shows the time-varying modularity index (*Q*) for Face and Animal stimuli observed at each threshold.

Analysis of the modularity index *Q* also showed that, in response to a face, the network modularity is reduced (5 %: *t*_max_ = 604, *p*-value < 0.05; 10 %: *t*_max_ = 559, p-value < 0.05; 20 %: *t*_max_ = 550, *p*-value < 0.05), suggesting a topology characterized by progressively integrated rather than segregated organization (Figure 5B).

## 4. Discussion

The present study sought to understand the brain dynamics subtending face processing from a network science perspective. We presented participants with a series of static images displaying either a face or an animal that the participants had to distinguish from a similar distractor and recorded their cortical responses using EEG. Applying several metrics obtained from graph-theory, we characterized these cortical responses in terms of event-related topological changes of the timevarying network derived from the alpha phase-locking value computed between each possible pair of cortical sites.

Since a typical electrophysiological signature of face processing is represented by the occipito-temporal N170 component (Rossion, 2014), we first predicted to observe that nodes in these regions would show an increased centrality, peaking around 170 milliseconds after stimulus onset.

Permutation-based analyses provided a strong support for this prediction, revealing that the sensors located over occipital and temporo-parietal areas (most prominently in the right hemisphere) are the ones characterized by the greatest centrality when the network is prompted with a face compared to its activity elicited by the view of an animal (Figure 2). Moreover, the timecourse of the centrality metrics shows that the difference in the network response to these two types of stimuli starts very early and peaks in the expected time frame (Figure 3). It is important to stress that these effects were observed for each of the three different metrics used to assess node centrality (degree, eigenvector centrality and PageRank centrality) and were invariant to the levels of sparsity of the connectivity matrices. Thus, the observed dynamics should be considered as an expression of inherent properties of the network activity triggered by a face.

Several models of the face-processing network posit that its core regions are the inferior occipital gyrus, the fusiform gyrus and the superior temporal sulcus (Haxby & Gobbini, 2012; Haxby et al., 2000; Ishai, 2008; Rossion et al., 2003), which are regions dedicated to the analysis of the visual features that define a face as an holistic percept (Haxby & Gobbini, 2012). Furthermore, these regions have been consistently found as the source of the N170 potential observed with EEG (Eimer, 2012; Jacques et al., 2019; Tadel et al., 2019). The present results bolster the claim for a distributed processing mechanism for faces (Haxby & Gobbini, 2012) that is centered around these regions, providing a quantitative measure of their involvement within the whole brain network. Indeed, our analysis show that the number of functional pathways involving occipital and temporoparietal nodes is larger when the stimulus to process is a face compared to a non-face stimulus. This is in line with recent studies that investigated using MRI, both functional (Zhen, Fang, & Liu, 2013) and structural (Pyles, Verstynen, Schneider, & Tarr, 2013) connectivity subtending face processing. Furthermore, the time-resolution granted by EEG allows to extend this evidence suggesting that the role of these regions is not static. Their role as functional hubs within the network is subtended by a dynamic adjustment of their influence, which begins as early as ~100 msec. Indeed, we observed that the centrality of these nodes is larger for faces than for non-face stimuli from the initial stages of processing, in contrast with classical ERP studies suggesting that face-sensitive neural activity arises only in the N170 component. In this study, the analysis of the cortical activity in terms of event-related network changes reveals an earlier primacy of face over non-face stimuli, in line with a very recent study which, combining EEG and fMRI, showed that the coding of facial expressions from occipital to the ventral temporal cortex starts in this time frame (Muukkonen, Ölander, Numminen, & Salmela, 2020). Furthermore, this result supports another recent study showing that face-sensitive neural activity can be detected already in the P1 component (Tanaka, 2018). Then, as stimulus processing continues to the next stages, we can observe a marked increase in the centrality of posterior hubs peaking around 200 milliseconds, likely reflecting the activity within the core system concerned with the coding of those visual features that uniquely distinguish a face. Finally, the centrality of these nodes drops around 300 milliseconds, marking the end of the neural computation needed for extracting the *faceness* from the stimulus (Rossion, 2014).

To better characterize the relationship between event-related potentials and event-related network changes we analyzed the statistical influence of the latter on the former using Granger causality, hypothesizing that the centrality scores can be associated with the electrical potentials collected at each node. Results supported our hypothesis, revealing that right temporo-parietal sensors centrality (for each metric considered) was indeed predictive of the amplitude of the ERP measured at these sites (Figure 4). This result showed that the larger the information exchange involving these nodes, the larger their recorded activity would be. Specifically, the increased activity of a neural source expressed in the ERP amplitudes might be considered the result of the increased connectivity that is dynamically established among all the neural sources involved in the face processing task, providing a causal (in the statistical Granger sense) link between the state of the network and the ERP response. Nevertheless, some caution is advisable when interpreting this result, since the effect is no longer observed when constraining the connectivity matrix to a very stringent level of sparsity. A possible explanation for this inconsistency is that centrality estimated from an overly sparse connectivity matrix, in which the overall number of possible links that a node can establish is necessarily low, magnifies large-scale dynamics at the cost of weaker connections which are nonetheless important for understanding the relationship between network state and local neuronal activity (Garrison, Scheinost, Finn, Shen, & Constable, 2015; Van Dijk et al., 2010).

Finally, we were interested in understanding how face processing is related to the global network topology, to test if the complex features defining a face are processed in discrete and segregated modules (Kanwisher & Barton, 2011; Kanwisher & Yovel, 2006) or instead the mechanism relies on a distributed but integrated system (Haxby & Gobbini, 2012; Schrouff et al., 2020). Our results support the latter hypothesis by showing that network topology is dynamically shaped during face processing toward a small-world organization characterized by strong integration (Figure 5). The analysis of small-worldness reveals indeed that the functional network exhibits a small-world topology (−0.5 < ω < 0.5) throughout the whole task for both faces and animals processing. This extends the evidence that the brain network has this inherent topological organization, mostly observed at rest, to an active condition. Moreover, we observed that ω progressively approaches 0 during the ongoing elaboration of both stimuli. This suggests a dynamical reconfiguration of the functional pathways mediating the visual computation that reaches a climax in the time-frame in which the N170 is typically observed. In line with the electrophysiological evidence showing that N170 reflects not a face-specific but rather a face-sensitive activity, here we show that network’s small-worldness follows the same principle. Nevertheless, the observation that ω is maximally reduced in response to the presentation of a face, suggests that its exhaustive visual processing requires a network that is dynamically reconfiguring its structure to have strong small-world architecture maximizing its efficiency (Bassett & Bullmore, 2006). Additionally, we observed that the modularity index *Q* is also characterized by a dynamic modulation, but only during face processing. Specifically, we observed that the network modularity is progressively reduced when a face is presented, again reaching its climax in the N170 time frame. Recent fMRI studies observed that the resting state functional connectome undergoes rhythmic fluctuations from states of high to low modularity, suggesting that these fluctuations reflect time windows in which the brain transitions from a state of high segregation to a state of high integration in which information is more freely exchanged across nodes (Betzel et al., 2016; Liao, Cao, Xia, & He, 2017; Sporns & Betzel, 2016). This evidence has been further extended to a condition of active processing (Di et al., 2013; Shine et al., 2016), revealing that the dynamic interplay between states of segregation and integration underscores behavioral performance in a simple attentional task, which is maximal when the network is in a state of strong integration (Shine et al., 2016). Here we further extend this evidence, showing that modularity transitions can be observed on the fast temporal scale of EEG, and that these dynamic changes subtend the processing of a complex visual stimulus like a face. The large reduction in modularity observed during face presentation, reveals that its recognition is mediated by a network configuration that maximizes the fast and efficient integration of information at the expense of a segregated subdivision. Thus, this result provide support for theoretical accounts positing that the face network relies on a distributed mechanism (Haxby & Gobbini, 2012) rather than a segregated architecture that operates on functionally separated pathways. Furthermore, they provide additional support for the evidence suggesting a strong integration among the several components of the face network in tasks that require an explicit judgment based on affective features (Fairhall & Ishai, 2007; Summerfield et al., 2006).

## 5. Conclusions

Altogether, the present results show that the efficient processing of a facial expression seems to rely not on the isolated activity of functional specialized modules acting in parallel on different visual features (Kanwisher & Yovel, 2006). Rather it results from the disruption of this segregation, which allows the emergence of a collective activity within the nodes comprising the network (Behrmann & Plaut, 2013). These need to be arranged in a small-world topology that maximizes the information exchange (Bassett & Bullmore, 2006; Telesford et al., 2011), with few highly connected hubs, likely corresponding to the core part of the face-perception network (Ishai, 2008; Jacques et al., 2019), whose influence dynamically increases as the computation progress. Finally, it is important to highlight that this study does not come without limitations. First, it is important to acknowledge that the limited scalp coverage prevented a finer spatial exploration by means of source reconstruction algorithms. Nevertheless, the use of CSD transform of the EEG data before connectivity analysis definitely mitigated the problem of volume conduction that can arise when investigating scalp connectivity (Kayser & Tenke, 2015). Additionally, the strong accordance with the topography observed in this study with previous EEG studies allows confidence in our conclusions. Second, it is important to highlight that in this study we focused only on the connectivity in the alpha band, due to its coordinating role across brain rhythms (Bahramisharif et al., 2013; van Driel et al., 2014). Nevertheless, the core of the visual computation lies in the cross-communication occurring at multiple oscillatory rhythms. Future studies should aim to extend our findings investigating how the present patterns generalize to other oscillatory scales and, more importantly, what are the cross-spectral functional connectivity patterns (Iandolo et al., 2020) subtending this complex network activity.

## CRediT authorship contribution statement

**Antonio Maffei:** *Conceptualization, Methodology, Formal analysis, Visualization, Writing - Original Draft*

**Paola Sessa:** *Conceptualization, Investigation, Data Curation, Supervision, Writing - Review & Editing*

## Ethics approval

The present research received approval by the Ethical Committee for psychological research of the University of Padova (Protocol number: 1986).

